# Genome-wide Association Study of Pediatric Obsessive-Compulsive Traits: Shared Genetic Risk between Traits and Disorder

**DOI:** 10.1101/858241

**Authors:** Christie L. Burton, Mathieu Lemire, Bowei Xiao, Elizabeth C. Corfield, Lauren Erdman, Janita Bralten, Geert Poelmans, Dongmei Yu, S-M Shaheen, Tara Goodale, OCD Working Group of the Psychiatric Genomics Consortium, Noam Soreni, Gregory L. Hanna, Kate D. Fitzgerald, David Rosenberg, Gerry Nestadt, Andrew D. Paterson, Lisa Strug, Russell J. Schachar, Jennifer Crosbie, Paul D. Arnold

**Author notes:** **Corresponding Author:** Christie L. Burton, PhD, Research Associate, The Hospital for Sick Children (Neurosciences and Mental Health/Psychiatry), 555 University Ave, 4289B, Toronto, ON M5G 1X8, ex 202346/Fax.

## Abstract

**Objective:** To identify genetic variants associated with obsessive-compulsive (OC) traits and test for sharing of genetic risks between OC traits and obsessive-compulsive disorder (OCD).

**Methods:** We conducted a genome-wide association analysis of OC traits using the Toronto Obsessive-Compulsive Scale (TOCS) in 5018 unrelated Caucasian children and adolescents from the community (Spit for Science sample). We tested the hypothesis that genetic variants associated with OC traits from the community would be associated with clinical OCD using a meta-analysis of three OCD case-controls samples (cases=3384, controls=8363). Shared genetic risk was examined between OC traits and OCD in the respective samples using polygenic risk score and genetic correlation analyses.

**Results:** A locus tagged by rs7856850 in an intron of *PTPRD* (protein tyrosine phosphatase δ) was significantly associated with OC traits at the genome-wide significance level (*p*=2.48×10^−8^). The rs7856850 locus was also associated with OCD in a meta-analysis of three independent OCD case/control genome-wide datasets (*p*=0.0069). Polygenic risk scores derived from OC traits were significantly associated with OCD in a sample of childhood-onset OCD and vice versa (*p*’s<0.01). OC traits were highly but not significantly genetically correlated with OCD (*r*_*g*_=0.83, *p*=0.07).

**Conclusions:** We report the first validated genome-wide significant variant for OC traits. OC traits measured in the community sample shared genetic risk with OCD case/control status. Our results demonstrate the importance of the type of measure used to measure traits as well as the feasibility and power of using trait-based approaches in community samples for genetic discovery.

---

Obsessive-compulsive disorder (OCD) is a common (1-2% prevalence)(1) psychiatric disorder characterized by intrusive, recurrent thoughts and repeated, ritualized behaviors. Up to 50% of OCD cases have a childhood-onset (before the age of 18)(2), which is more heritable than adult-onset OCD (3). Two genome-wide association studies (GWAS) in clinical samples with mixed ages of OCD-onset and a meta-analysis of these studies did not identify genome-wide significant findings (4–6). Top hits from previous GWAS include SNPs within *DLGAP1, BTBD3, GRID2* and one close to *PTPRD*. Using obsessive-compulsive (OC) symptoms rather than a clinical diagnosis, a study of adult twins identified a genome-wide significant SNP in *MEF2B* (rs8100480)(7). However, this SNP was not replicated in an independent sample (5).

We conducted a GWAS of quantitative OC traits in a large pediatric, community-based sample: Spit for Science (8). We measured OC traits using the Toronto Obsessive-Compulsive Scale (TOCS - 8). This heritable measures (9) that includes negative scores that represent ‘strengths’ (e.g., never upset when their belongings are rearranged) and positive scores that represent ‘weaknesses’ (e.g., very upset when their belongings are rearranged). We reasoned that a strength-to-weakness format would generate scores with a more normal distribution in a community sample (8) than those observed with typical OCD scales and would therefore boost power of genetic discovery (10). Typical OCD trait measures generate J-shaped distributions because their format calls for ratings of symptoms from absence to presence (score of zero to a positive integer). A j-shaped distribution is especially likely when using typical OCD measures in a community sample where the prevalence of OC symptoms is low and most people would get scores of zero (11). This j-shaped distribution can be replicated with the TOCS by collapsing the ‘strengths’ (i.e., negative scores) into scores of zero (Figure 1). We tested the hypothesis the distribution of TOCS scores would boost power of genetic discovery (10) by running a GWAS with the collapsed TOCS measure as well as the full distribution. We characterized the genetic associations for TOCS by conducting gene-based analyses, examining brain expression quantitative trait loci (eQTLs) of top hits, estimating SNP-based heritability and genetic correlations of total OC trait scores with other medical/mental health disorders and traits. We also examined if the top hits from the previous GWAS of OC symptoms (7) replicated in our study. Finally, we tested the hypothesis that OC traits in the community share genetic risk with OCD by examining individual genetic variants, genetic correlations and polygenic risk between OC traits in Spit for Science and three independent OCD case/control samples.

**Figure 1:**
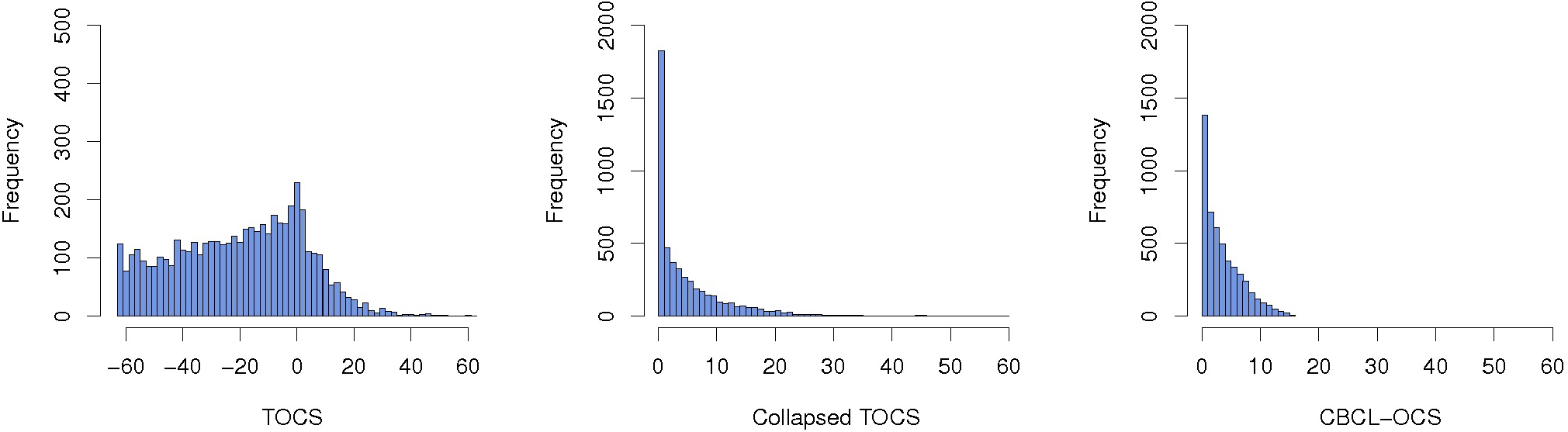
Distribution of OC Trait Measures. Histograms of a) TOCS total score, b) total score for collapsed TOCS items (all negative scores converted to zero for each item) and c) the Child Behavior Checklist – Obsessive-Compulsive Scale (CBCL-OCS). *n*=5018

## Materials and Methods

### OC Traits

#### Participants

The Spit for Science sample is described in detail elsewhere (8). Briefly, the sample included 15 880 participants with complete demographic, questionnaire and family information (mean age=11.1 years [SD 2.8]; 49.4% female) from the 17,263 youth (6-18 years of age) recruited at the Ontario Science Centre over 16 months. Informed consent, and assent where applicable, were obtained using a protocol approved by The Hospital for Sick Children Research Ethics Board. Participants provided a saliva sample in Oragene saliva kits (OG-500; DNA Genotek, Ottawa, Canada) for genetic analyses. See the supplement for details.

#### OC Trait Measure

We measured parent- and self-reported OC traits within the last 6 months using the TOCS, a 21-item questionnaire described previously (8,9). Each item was scored on a 7-point Likert scale ranging from −3 (‘far less often than others of the same age’) to +3 (‘far more often than others of the same age’). A score of zero was designated as an average amount of time compared to same-age peers. The TOCS total score was standardized into a z-score to account for age, sex and questionnaire respondent (parent or self). Details of z-score creation are described in the supplement. We tested the impact of the strength/weakness structure of the TOCS by re-scoring the TOCS to convert all negative scores for individual items to zero before summing scores (i.e., no scores less than 0 which collapsed the left side of the distribution). We also compared the TOCS to an additional OCD symptom measure with a j-shaped distribution: CBCL-OCS (12). Each of the eight CBCL-OCS items was scored on a scale of 0-2 (0 = not true; 1 = somewhat/sometimes true; and 2 = very/often true) and was summed to generate a total score (range: 0-16). This ‘collapsed’ TOCS total score, with a cluster of scores at zero, created a distribution similar to the CBCL-OCS (Figure 1)

#### Genetic Data

DNA was extracted manually from saliva using standard methods (see the supplement for additional details). We excluded any samples with concentrations <60ng/µl and insufficient quality based on agarose gels. We genotyped 5645 samples on the Illumina HumanCoreExome-12v1.0_B (HumanCore) and 192 samples on the Illumina HumanOmni1-Quad V1.0_B (Omni) bead chip arrays (Illumina, San Diego, CA, USA) at The Centre for Applied Genomics (Hospital for Sick Children, Toronto, CA). There were 538 448 markers on the HumanCore and 1 140 449 markers on the Omni array.

Quality control (QC) was conducted separately for each array using standard methods with PLINK v1.90 (13). Sample exclusion and selection criteria are described in the supplemental methods and Supplemental Figure S1. Imputation was performed separately for all platforms and sample sets, using Beagle v4.1 using the data from phase 3, version 5 of the 1000 Genomes project for reference (http://bochet.gcc.biostat.washington.edu/beagle/1000_Genomes_phase3_v5a/). We excluded individuals who were non-Caucasian based on principal component (PC) analysis and included only one participant from each family (inferred sibs or half-sibs, see supplement and Supplemental Figure S2).

#### Analyses

GWAS were conducted using R (v3.5.1). Our primary analysis tested if imputed dosage and standardized TOCS total score were associated using a linear regression model that included the top three PCs and genotyping array as covariates. We included SNPs with a minor allele frequency (MAF)>1%, allelic R^2^ imputation quality (AR2>0.3) and used the standard genome-wide threshold of *p*≤5×10^−8^. We also tested if any genome-wide significant variants from the analysis with the standardized scores were still significant using a non-standardized TOCS score. For these analyses, age, sex, respondent and their 2- and 3-way interactions were used as covariates in addition to the above (interactions were included to mimic the construction of the Z scores, which were calculated independently in age-, sex- and respondent-defined bins; see supplement).

In secondary analyses, we evaluated the association between SNPs and the collapsed TOCS score, and between SNPs and CBCL-OCS, using zero-inflated negative binomial likelihood ratio tests, using the function *zeroinfl* from the *R* package *pscl* (v1.5.2). This model was chosen because of the high proportion of zero scores that created a j-shaped distribution. The test is a mixture of two models: a negative binomial model, which contributes to zero and positive scores and a logit model, which contributes to possible inflation of zero scores (point mass at 0) compared to what a negative binomial model predicts. These analyses used non-standardized scores for the collapsed TOCS and CBCL-OCS so the model adjusted for the covariates and the association of SNP allele dosage with the OC trait scores is tested against the null of having no effects on both the logic part and the negative binomial part using likelihood ratio tests.

We subsequently used FUMA to conduct gene-based GWAS of the TOCS standardized total score with MAGMA using a Bonferonni correction for number of protein coding genes included (19 - fuma.ctglab.nl).

We tested each genome-wide significant variant for co-localization with brain eQTLs using LocusFocus (https://locusfocus.research.sickkids.ca/ (20). We examined the 14 GTEx sets from brain tissue types and examined SNPs within ±1 Mbp of each SNP.

We estimated SNP heritability using both GCTA (16) v1.91.2-beta (http://cnsgenomics.com/software/gcta/) with further exclusion of cousins and SNPs with AR2>0.9 and LDSC (v1.0.0, https://github.com/bulik/ldsc; 22) calculated from SNPs in HapMap3. We used LDSC (18) to examine the genetic correlation of TOCS total scores with the 850 phenotypes available on LD Hub (http://ldsc.broadinstitute.org/ldhub/).

Finally, we examined the top variants in the only previous GWAS of OC symptoms (only 20 loci reported; 7) in the results from the TOCS total score GWAS.

### OCD Case/Control

#### Participants

For validation analyses, we investigated three independent OCD case/control cohorts: 1) the International OCD Foundation Collaborative (IOCDF-GC) and OCD Collaborative Genetics Association Studies (OCGAS) meta-analysis - 6), 2) the Philadelphia Neurodevelopmental Cohort (PNC) from the Children’s Hospital of Philadelphia (CHOP - 24) and 3) the Michigan/Toronto OCD Imaging Genomics Study (20). See the supplement and Table 1 for sample sizes.

**Table 1:** Overview of Samples

| **OC Trait Sample** | **Phenotype** | **Age** | **Total Samples** | **Cases** | **Controls** |
| --- | --- | --- | --- | --- | --- |
| Spit for Science | TOCS, CBCL-OCS | 6-18 years | 5018 | n/a† | n/a |

| **All Replication** | **Phenotype** | **Age of Onset** |  |  |  |
| --- | --- | --- | --- | --- | --- |
| IOCDF/OCGAS | Clinician-Diagnosed (DSM-IV) | Child and adult | 9725 | 2688 | 7037 |
| CHOP | Case/control status based on GO-ASSESS symptoms | <18 years of age | 1775 | 406 | 1369 |
| Michigan/Toronto | Clinician-Diagnosed (DSM-IV) | <18 years of age | 480 | 275 | 205 |
| **Total Meta-Analysis** |  |  | **11980** | **3369** | **8611** |

† There were 62 individuals that had a parent- or self-reported diagnosis of OCD. Number of samples reflect samples included in analyses (i.e., after quality control analyses). IOCDF/OCGAS = International OCD Foundation Collaborative and OCD Collaborative Genetics Association Studies meta-analysis sample, CHOP = the Philadelphia Neurodevelopmental Cohort (PNC) from the Children's Hospital of Philadelphia, TOCS = Toronto Obsessive-Compulsive Scale, CBCL-OCS = Child Behavior Checklist – Obsessive-Compulsive Scale, DSM-IV = Diagnostic and Statistical Manual of Mental Disorders, 4<sup>th</sup> Edition

#### Analyses

GWAS summary stats from each OCD sample were meta-analyzed using fixed-effect inverse variance methods. We tested if the results from the gene-based GWAS for the TOCS total score replicated in the OCD samples by conducting the same genome-wide analyses as described above or in the supplement. Polygenic risk score (PRS) analyses were performed using LDpred v1.06 (21) (see supplement). First, we derived PRS for TOCS from the Spit for Science sample and tested their association with case/control status in the combined OCD cohorts (target sample: CHOP, Michigan/Toronto and a subset of the IOCDF-GC/OCGAS - see supplement). Second, we derived PRS from the combined OCD cohorts and tested their association with the standardized TOCS total score in the Spit for Science sample (target sample).

We examined the potential shared genetic risk between the Spit for Science and the meta-analyzed OCD samples using genetic correlations estimated with LDSC (18).

## Results

### OC Traits

We used 5018 participants for GWAS analyses after sample exclusion and selection (see supplement and Supplemental Figures S1/S2). In the primary analysis, rs7856850 in *PTPRD* was significantly associated with TOCS total scores at the genome-wide level (*p*=2.48×10^−8^, β=0.14, s.e.=0.025, *R*^2^=0.618%: Figure 2a, top hits listed in Supplemental Table S1). Several variants in this region that approached genome-wide significance were in linkage disequilibrium (LD) with rs7856850, which was genotyped on both the HumanCore and OMNI arrays (Figure 2B). The inflation factor λ was 1.008 while the intercept of LD score regression was 1.003 and not significantly different from 1 (s.e.=0.007, *p*=0.66; Figure 2C). rs7856850 was still associated with TOCS total scores using raw instead of standardized scores (*p*=2.75 × 10^−8^, *R*^2^ = 0.615%; data not shown).There was no eQTL data for the SNP in *PTPRD*, rs7856850 in LocusFocus (15) or in the most recent version of GTEX v8 (22).

**Figure 2:**
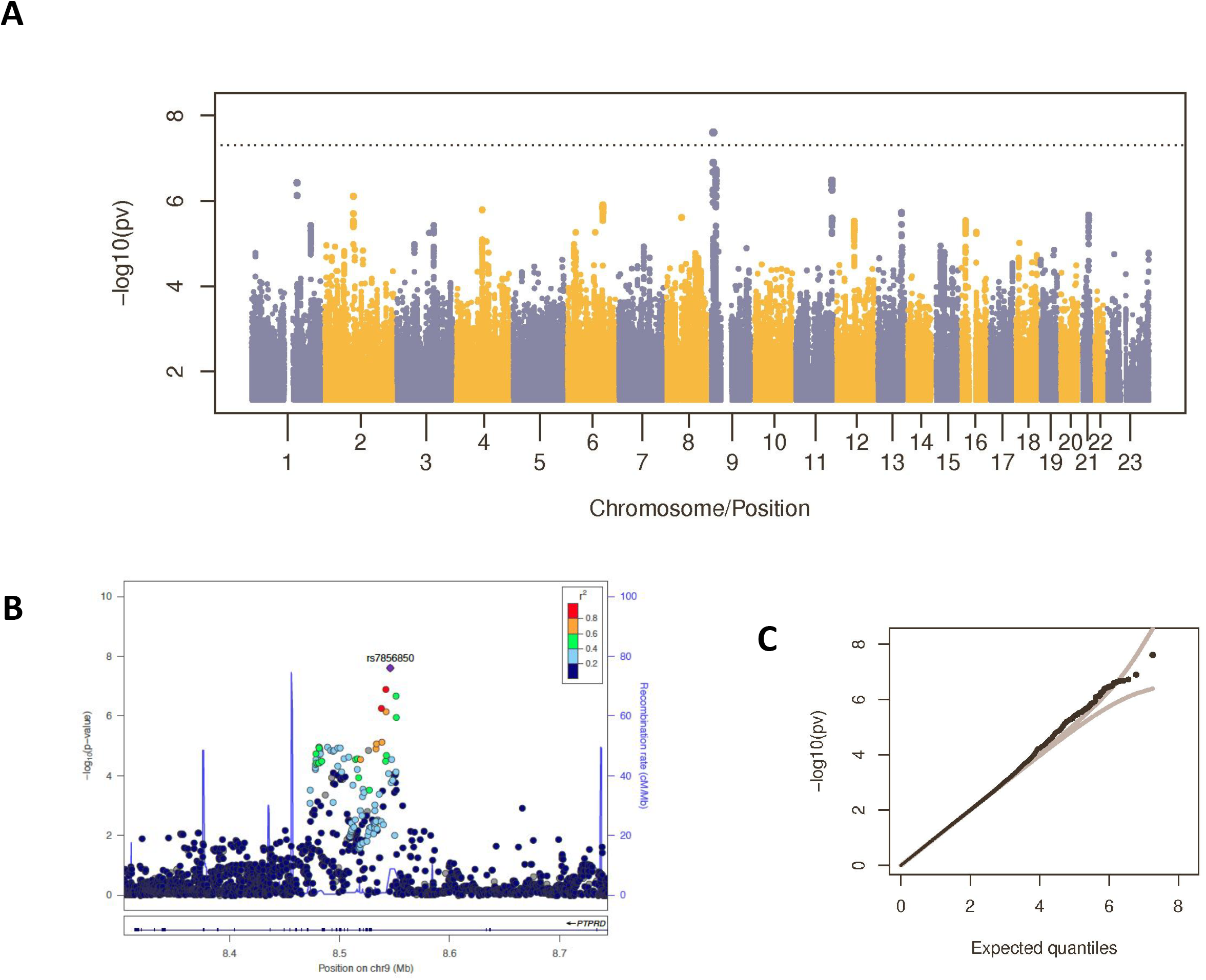
Genome-Wide Significant Locus in *PTPRD* Associated with OC Traits in Spit for Science. a) Manhattan plot for GWAS of the TOCS standardized total score. rs7856850 in one the introns of *PTPRD* surpassed the genome-wide threshold (*p*=5×10^−8^; grey line). b) Locus zoom plot for the genome-wide significant locus from the GWAS of the TOCS standardized total score. c). QQ Plot for the GWAS of the TOCS standardized total score. *n*=5018

When we analyzed the collapsed TOCS total score and the CBCL-OCS, the genome-wide significant locus for the TOCS total score rs7856850 was no longer genome-wide significant, although the remaining effect was in the same direction had the same direction of effect (*p*=0.00045 and *p*=0.025 respectively; see supplement for details). For both collapsed TOCS and the CBCL-OCS, the A allele was associated with both higher scores (collapsed TOCS: β=0.0735, s.e.=0.0292, CBCL-OCS: β=0.0465, s.e.=0.020) and lower proportion of zero scores (collapsed TOCS: β=-0.231, s.e.=0.104, CBCL-OCS: β=-0.126, s.e.=0.179).

A gene-based GWAS of the TOCS standardized total score using MAGMA on the FUMA platform identified four genome-wide significant genes (at a Bonferroni-corrected level *p*= 0.05/19363 protein coding genes=2.582×10^−6^): *SH3GL2* (*p*=1.535×10^−7^, *z*=5.12); *PDXDC1* (*p*=3.95×10^−7^, *z*=4.94); RIMBP2 (*p*=1.513×10^−6^, *z*=4.70) and RRN3 (*p*=1.56×10^−6^, *z*=4.66; see Supplemental Figure S3). PDXDC1 and RRN3 have overlapping coding regions.

The heritability of the TOCS total score was *h*^2^=0.068 (s.e.=0.052, *p*=0.19) using GCTA and *h*^2^=0.073 (s.e.=0.064; *p*=0.25) using LDSC when the intercept was constrained to 1. TOCS total score was not significantly associated with any phenotypes on LD Hub (see supplement).

One of the top-ranked SNPs from a previous GWAS of OC symptoms (7) was nominally associated with TOCS total scores in the Spit for Science sample with the same direction of effect (rs60588302, *p*=0.025). This SNP is in the same region as our top hit (9p24.1), but not in LD (*r*^2^=0.004, D’=0.517). Another 16 of the reported top hits in den Braber (7), including a variant in *MEF2BNB* (rs8100480) that was genome-wide significant in their sample, had effects in the same direction but were not significantly associated in the current sample (Supplemental Table S2).

### OCD Case/Control

Following standard QC and sample exclusion where applicable (see supplement), we had a total of 3369 cases and 8611 controls in our validation samples (Table 1). We tested if the genome-wide SNP associated with TOCS total scores in Spit for Science were also associated with OCD in the meta-analyzed cohorts. rs7856850 was associated with increased odds of being an OCD case (*p*=0.0069, OR=1.104 per A allele [95% confidence limit 1.03-1.19], Figure 3, Supplemental Figure S4). A gene-based GWAS by MAGMA using FUMA did not identify any genome-wide significant genes. Therefore, none of the genes identified in the gene-based GWAS of OC traits were replicated (see supplement). The top three genes in the analysis were *GRID2* (*p*=7.52×10^−6^, *z*=4.33), *SDCBP2* (*p*=9.05×10^−6^, *z*=4.29) and *DLGAP1* (*p*=9.05×10^−6^, *z*=4.29; Supplemental Figure S5).

**Figure 3:**
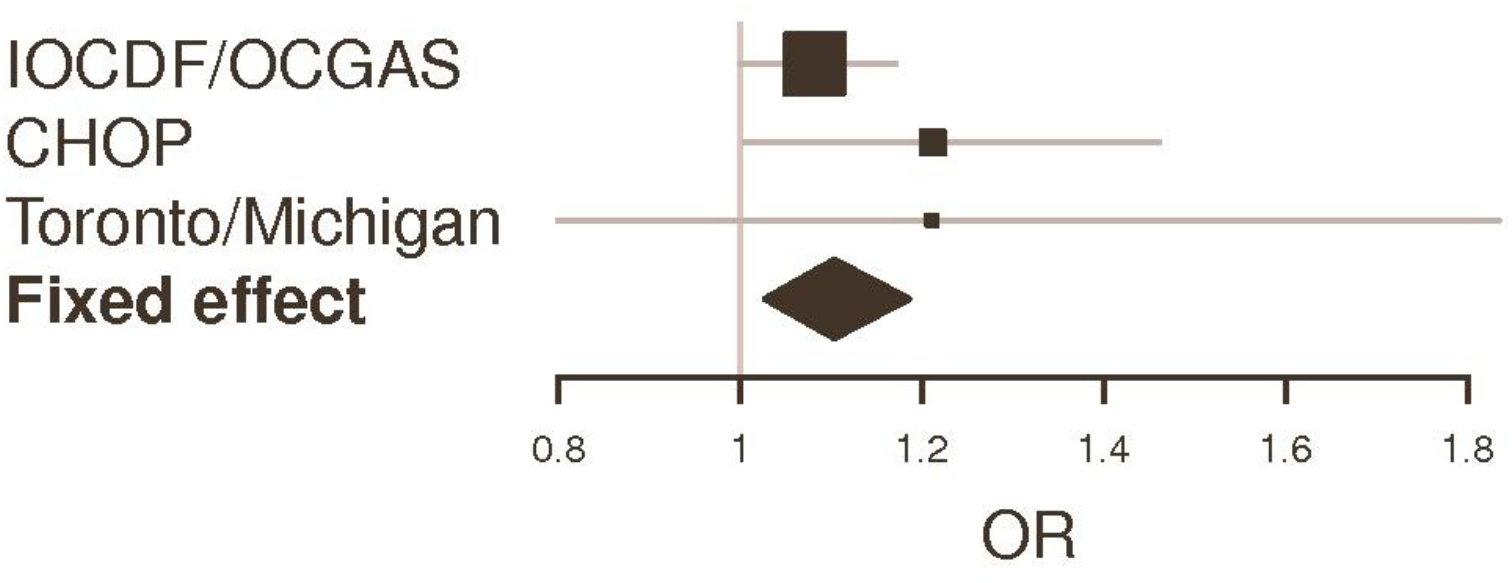
Validation of Locus in PTPRD in OCD Samples. Forrest plot of genome-wide significant variant (rs7856850) across all the replication samples and sub-samples: 1) IOCDF/OCGAS (International Obsessive-Compulsive Disorder Foundation Collaborative and OCD Collaborative Genetics Association Studies samples), 2) CHOP (Philadelphia Neurodevelopmental Cohort (PNC) from the Children’s Hospital of Philadelphia and 3) Michigan/Toronto OCD Imaging Genomics Study. OR = Odds Ratio

Figure 4a shows that PRS calculated for TOCS total scores was significantly associated with increased odds of being a case in the meta-analyzed OCD samples (Nagelkerke’s pseudo *r*^2^=0.277%, *p*=0.0045 at ρ=0.003). Figure 4b shows that PRS constructed from the OCD sample were significantly associated with TOCS total scores in Spit for Science (*r*^2^=0.24%; *p*=0.00057 at ρ=0.1).

**Figure 4:**
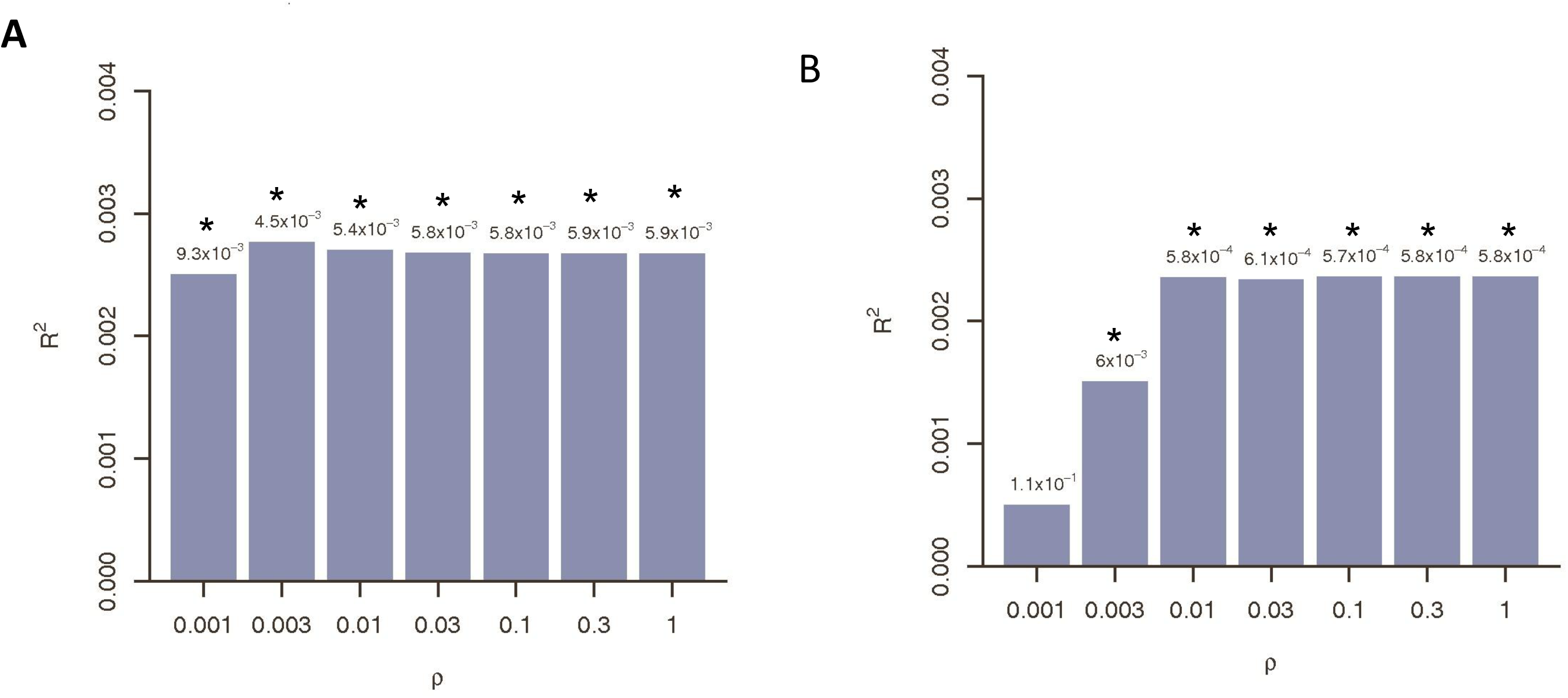
OC Traits in the Community and OCD Share Polygenic Risk. a) Variance explained (R^2^) in OCD case/control status in replication samples by polygenic risk for OC traits from Spit for Science b) variance explained in OC traits in Spit for Science sample by polygenic risk for OCD from replication samples across a range of prior proportion of causal variants (ρ). Analyses conducted using LDpred. * = p < 0.01

The genetic correlation between standardized TOCS total scores and OCD meta-analysis was *r*_*g*_=0.825 (s.e.=0.428; *p*=0.073) when intercepts are constrained to 1.

## Discussion

Using a trait-based approach in a community sample, we identified a genome-wide significant variant associated with OC traits (rs7856850) that was also associated with OCD case/control status. Polygenic risk and genetic correlation findings showed sharing of genetic risks between OC traits in the community and OCD case/control status in independent samples.

The genome-wide significant variant (rs7856850) associated with OC traits is in an intron of the consensus transcript of *PTPRD* that codes for protein tyrosine phosphatase δ. No eQTLs have been calculated yet for rs7856850 (GTEx V8; 20). This variant was validated in a meta-analysis of three independent OCD cohorts making it the first variant associated with OC traits and OCD. The small size of the meta-analyzed OCD sample likely precluded finding genome-wide hits in the meta-analysis. We did not validate the genes associated with OC traits using MAGMA in the OCD sample. Previous GWAS of OCD symptoms or diagnosis identified variants that approached significance in the region around *PTPRD*. However, those variants were independent of the locus found in our study (4,7). These observations support a possible role of the 9p24.1 region in OCD. The 9p region is also the location of one of the strongest linkage peaks in earlier genome-wide linkage studies of pediatric OCD (23,24). Rare CNVs in *PTPRD* have been identified in cases with OCD (20) and ADHD (25). SNPs in *PTPRD* were genome-wise significantly associated with ASD (26), restless legs syndrome (27), and self-reported mood instability (28). *Ptprd*-deficient mice show learning deficits and altered long-term potentiation magnitudes in hippocampal synapses (29). *PTPRD* is expressed highly in the brain compared to non-brain tissues, especially in myelinating axons and growth cones (30,31) in the prenatal cerebellum (32). The presynaptically located *PTPRD* is involved in axon outgrowth and guidance (33) and interacts with postsynaptic proteins such as Slitrk-2, interleukin-1 receptor and TrK to mediate synapse adhesion and organization in mice (34,35) and the development of excitatory and inhibitory synapses (36). Members of the Slitrk and interleukin protein families have been associated with OC behaviors in humans and mice (37,38).

Our results show that OC traits in the community share genetic risk with OCD. Polygenic risk for OC traits was associated with OCD case/control status and vice versa. OC traits and OCD case/control status also were substantially, but not significantly, genetically correlated. This estimate is higher than reported in a recent study (*r*_*g*_=0.42, *p*=0.095; 50). Lack of power is the most likely explanation for the absence of significant results. Previous studies on other psychiatric disorders report shared genetic risk between traits and diagnoses, with polygenic risk and genetic correlations similar to what we report for OC traits and OCD case/control status (28,40,41). The shared genetic risk between OC traits and OCD supports the hypothesis that an OCD diagnosis could represent the high extreme of OC traits that are widely distributed in the general population. One implication of this finding is that population-based samples with quantitative trait measures can serve as a powerful complementary approach to case/control studies to accelerate gene discovery in psychiatric genetics.

SNP-based heritability for OC traits in the current sample was not significant in line with previous studies. Previous research reports lower SNP-based heritability for self-reported OC symptoms (0.058) (39) than for clinical OCD (0.28-.37)(6,42). A similar trend for lower trait vs. diagnosis SNP heritability has been observed for ADHD (40,43). The reason for the disparity in SNP heritability between traits and diagnosis is unclear. One possible explanation could be differences in the informant (parent/self vs. teacher or clinician) as shown previously for ADHD (43). Regardless of a non-significant SNP heritability for OC traits from our sample, we still identified and validated a genome-wide significant variant.

The TOCS scale is similar to existing OC trait/symptom measures in item content, but is unlike existing scales in that it measures OC traits from ‘strengths’ to ‘weaknesses’. As a result, the distribution of the total score is closer to a normal distribution than the j-shaped distributions typically observed with most symptom-based scales that rate behaviors from zero to a positive integer (9 - e.g., not at all to quite a lot). Our results indicate that the distribution of the OC trait measure impacts power to identify genome-wide significant associations. A ‘strengths’ to ‘weaknesses’ measure identified a genome-wide significant association. However, when we collapsed the ‘strength’ end of the TOCS distribution to zero the significance of this variant was substantially reduced to below genome-wide significance, although the effect was in the same direction. The same effect was identified using another OC measure that generates a j-shaped distribution: CBCL-OCS. One implication of our results is that there is genetic information in the ‘strengths’ end of the distribution captured by the TOCS. This information would be lost in scales that only measure ‘weaknesses’, particularly in community samples where base rates of OC traits are relatively low. Trait-based scales that capture ‘strengths’ and ‘weaknesses’ and have a less skewed distribution could improve power to identify genome-wide hits and variants associated with disorders, especially in population samples where the prevalence of clinically significant OC symptoms is relatively low.

## Conclusions

We identified the first genome-wide significant variant for OC traits that was also associated with OCD case status. Power to detect a genome-wide association was impacted by the distribution of the OC trait measure. OC traits and OCD share genetic risks supporting the hypothesis that OCD represents the extreme end of widely distributed OC traits in the population. Trait-based approaches in community samples using measures that capture the whole distribution of traits is a powerful and rapid complement to case/control GWAS designs to help drive genetic discovery in psychiatry.

## Supporting information

Supplemental Methods and Results

## Acknowledgements

This study was funded by the Canadian Institutes of Health Research (PDA: MOP-106573; RJS: MOP– 93696). PDA is supported by the Alberta Innovates Translational Health Chair in Child and Youth Mental Health. The OCD Collaborative Genetics Association Study (OCGAS) was funded by the following NIMH Grant Numbers: MH071507 (GN), MH079489 (DAG), MH079487 (JM), MH079488 (AF) and MH079494 (JK). Yao Shugart and Wei Guo were also supported by the Intramural Research Program of the NIMH (MH002930-06). The IOCDF-GC was supported by a grant from the David Judah Foundation (a private, nonindustry-related foundation established by a family affected by OCD), MH079489 (DLP), MH073250 (DLP), S40024 (JMS), MH 085057 (JMS), and MH087748 (CAM). Support for the collection of the data for Philadelphia Neurodevelopment Cohort (PNC) was provided by grant RC2MH089983 awarded to Raquel Gur and RC2MH089924 awarded to Hakon Hakonarson. All subjects were recruited through the Center for Applied Genomics at The Children’s Hospital in Philadelphia. The Michigan/Toronto sample was supported by the Ontario Brain Institute – Province of Ontario Neurodevelopmental Disorders (POND) Network (grant number: IDP-PND-2018; awarded to RJS, JC and PDA) and the National Institutes of Mental Health (MH101493 awarded to GH and PDA; MH085321 awarded to GH, DR and PDA). The authors wish to thank Tara Paton, Chao Lu and Jo-Anne Herbrick of The Centre for Applied Genomics, The Hospital for Sick Children, Toronto, Canada for assistance with genotyping. We gratefully acknowledge all the studies and databases that made GWAS summary data available: ADIPOGen (Adiponectin genetics consortium), C4D (Coronary Artery Disease Genetics Consortium), CARDIoGRAM (Coronary ARtery DIsease Genome wide Replication and Meta-analysis), CKDGen (Chronic Kidney Disease Genetics consortium), dbGAP (database of Genotypes and Phenotypes), DIAGRAM (DIAbetes Genetics Replication And Meta-analysis), ENIGMA (Enhancing Neuro Imaging Genetics through Meta Analysis), EAGLE (EArly Genetics & Lifecourse Epidemiology Eczema Consortium, excluding 23andMe), EGG (Early Growth Genetics Consortium), GABRIEL (A Multidisciplinary Study to Identify the Genetic and Environmental Causes of Asthma in the European Community), GCAN (Genetic Consortium for Anorexia Nervosa), GEFOS (GEnetic Factors for OSteoporosis Consortium), GIANT (Genetic Investigation of ANthropometric Traits), GIS (Genetics of Iron Status consortium), GLGC (Global Lipids Genetics Consortium), GPC (Genetics of Personality Consortium), GUGC (Global Urate and Gout consortium), HaemGen (haemotological and platelet traits genetics consortium), HRgene (Heart Rate consortium), IIBDGC (International Inflammatory Bowel Disease Genetics Consortium), ILCCO (International Lung Cancer Consortium), IMSGC (International Multiple Sclerosis Genetic Consortium), MAGIC (Meta-Analyses of Glucose and Insulin-related traits Consortium), MESA (Multi-Ethnic Study of Atherosclerosis), PGC (Psychiatric Genomics Consortium), Project MinE consortium, ReproGen (Reproductive Genetics Consortium), SSGAC (Social Science Genetics Association Consortium) and TAG (Tobacco and Genetics Consortium), TRICL (Transdisciplinary Research in Cancer of the Lung consortium), UK Biobank. We gratefully acknowledge the contributions of Alkes Price (the systemic lupus erythematosus GWAS and primary biliary cirrhosis GWAS) and Johannes Kettunen (lipids metabolites GWAS).

