## Supplemental Methods and Results for "Genome-wide Association Study of Pediatric Obsessive-Compulsive Traits: Shared Genetic Risk between Traits and Disorder"

**Supplemental Methods and Materials**

**OC Traits**

**Participants**

We collected behavioural information about the participants from themselves if they were deemed capable of self-reporting (18.6%; typically 12 years of age or older) or from their parents (81.4%). Ethnicity was estimated using a self-report questionnaire and confirmed using genetic data (see below). We collected information about whether the participant had ever received a diagnosis of, or had been treated, for OCD. The sample was highly enriched for siblings with 51.6% of the sample (*n*=8190) having at least one sibling who also participated in the study (total number of families = 3816).

**Z-Score Estimation**

We created standardized z-scores that accounted for age, sex, and respondent-type (parent or self), which were all associated with the TOCS total score (*p* < 0.05). In order to eliminate ties when estimating z-scores, the modified total scores were modeled separately by respondent, with age and sex as covariates, treating family as a random effect, and residual scores were used to calculate the z-scores. Children and youth were divided into thirty groups according to respondent, gender and age. For parent respondents, groups included every integer year of age from 6-15. For self-respondents, integer year age groups were created for ages 13-17. A normally distributed quantitative score was assigned to each subject by sorting the residuals and substituting a z-score corresponding to their empirical percentile within each group. Z-scores were assigned within each of the thirty groups separately so that the distribution of scores would be comparable across age, gender, and respondent.

**Preparation of Genetic Samples & Genotyping**

To precipitate any possible carbohydrates in the sample, we centrifuged the samples for an additional 10 min at 10000 RPM and removed any formed pellet from the sample. DNA was quantified using the Quanti-iT, Pico Green® dsDNA kit from Invitrogen (Thermo Fisher Scientific) and samples with concentrations < 60ng/µl were excluded (6.5% of all extracted cases). DNA was subsequently aliquoted and stored at -80°C. Prior to conducting the micro-arrays, DNA quality was verified using agarose gels and 98.5% of samples had sufficient DNA quality.

On each 96 well plate, we also genotyped an individual from a Caucasian HapMap trio (CEPH) as a quality control (1) with each sample genotyped approximately 20 times. Genotypes were called using GenomeStudio (Illumina, San Diego, CA, USA) separately for the HumanCoreExome (GenomeStudio v 1.9.4) and HumanOmni1 samples (GenomeStudio v2009.2).

**Genotyping QC and Selection of Participants for Genetic Analysis**

SNP position and annotation information were based on NCBI36 for Omni and on Genome Reference Consortium 37 (GRCh37) for HumanCore. Samples were excluded if their call rate was below 97%, a heterozygosity rate of 6 times the interquartile range from the closest quartile and/or their predicted and reported sex were mismatched. SNPs were excluded if they had call rates below 97%, they deviated from the rules of Hardy-Weinberg equilibrium at an FDR <1% (based on a set of homogeneous samples in terms of ancestry) and/or were duplicates of other SNPs, based on position and alleles (only the SNP with the highest call rate was retained). Nine participants were successfully genotyped on both platforms and we only kept data from the HumanCoreExome array. Samples were also excluded from statistical analysis if they did not have four grandparents of reported Caucasian descent, had a sex aneuploidy based on copy number analysis, did not have a standardized TOCS total score or had a parent- or self- reported diagnosis of autism spectrum disorder (ASD). Participants with ASD were excluded because the TOCS queries some behaviours that are common in ASD and we wanted to reduce the chance of phenocopies in our sample (2). Concordance of the HapMap trio samples genotyped on each HumanCore plate, were also >99.99%.

**Imputation**

A/T and C/G genotyped SNPs were removed prior to imputation. Allele coding on the X chromosome were coded as 0,1,2 for females and 0,2 for males.

**Ethnicity and Principal component calculation**

Figure S1 outlines the number of samples removed during QC. First, we excluded participants that did not have four grandparents of reported Caucasian descent. Next, principal components (PCs) were calculated from a set of autosomal, bi-allelic ancestry informative markers (AIM), calculated from samples from phase 3 of the 1000 Genomes project. We first pruned SNPs for linkage disequilibrium (*r^2^*<0.2 in 1500 kbp windows). Then, for each continental population, the top 1% SNPs with the largest frequency differences between that population and all others were retained. We ignored SNPs in the intervals chr8:7000000-13000000 [hg19] (8p23 inversion) and chr6:25000000-34000000 (MHC).

Participants’ AIMs were extracted from the imputed data sets, as long as their imputation quality was AR2>0.8. Hard genotype calls were used. To identify outliers with respect to ancestry, data from participants were combined with samples from the 1000 Genomes project. PCs were calculated using plink v1.90, and we excluded outliers in any of the first 3 principal components calculated from ancestry informative markers and from combining the participants with samples from phase 3 of the 1000 genomes project (see below; see Figure S2. Once ancestry outliers were removed, we recomputed PCs without 1000 Genomes samples.

**Relatedness**

The set of AIM SNPs (average observed heterozygosity of 0.41) was used to assess relatedness among the participants, using the “genome” option of plink. The estimated proportion π of autosomal genome identical by descent was inspected in pairs of participants. Networks of related participants with estimated pairwise π > 0.18 (inferred half-sibs or closer) was constructed. For Spit for Science samples, one participant from each network was retained for GWAS analysis, based on the highest standardized TOCS value. For the case-control studies, selection was based on participants being a case and/or older.

**OCD Case/Control**

**Participants**

*OCD Cohort 1: Meta-Analysis of the International Obsessive-Compulsive Disorder Foundation Collaborative (IOCDF-GC) and OCD Collaborative Genetics Association Studies (OCGAS) Samples.* This meta-analysis of two published GWASs of OCD in European Caucasians is described in detail elsewhere (3). The study consisted of 2688 patients with OCD based on DSM-IV criteria and 7037 genomically matched controls. Summary statistics from the IOCDF-GC/OCGAS GWAS were downloaded from <https://www.med.unc.edu/pgc/results-and-downloads/ocd/> (file ocd_aug2017.gz). Individual level data was accessed through dbGaP accession number phs000092.v1.p.

*OCD Cohort 2: Philadelphia Neurodevelopmental Cohort (PNC) from the Children’s Hospital of Philadelphia (CHOP).* The PNC sample is described in detail elsewhere (e.g., 4) and the data were obtained under approval from dbGaP phs00607.v2.p2. Briefly, the PNC sample is comprised of 9428 participants aged 8-21 years recruited from over 50 000 children genotyped from a blood sample by the Center of Applied Genomics after coming to CHOP or a CHOP-affiliated clinic for pediatric care. Participants were recruited randomly after the sample was stratified by age, sex and ethnicity (4). Participants completed a computerized structured screener based on the Kiddie-Schedule for Affective Disorders and Schizophrenia (K-SADS) called GO-ASSESS (5,6). The GO-ASSESS section related to OCD asked about the lifetime presence of any obsessive or compulsive symptoms as well as the severity, level of impairment and age of onset of symptoms.

In our validation analyses, we included individuals genotyped on the Illumina Human610-Quadv1_B BeadChip array who self-reported as European Caucasian. The samples released were all previously genotyped by the Center for Applied Genomics at The Children's Hospital of Philadelphia (7). QC was conducted as described for the Spit for Science sample. Participants were categorized into two groups: 1) participants with at least one OC symptom and reported impairment from the symptom(s) were considered cases (*n*=421), and 2) participants with no OC symptoms and reported impairment were considered controls (*n*=1441). We tested the association between imputed genotypes and case/control status using logistic regression controlling for age, sex and three principal components.

*OCD Cohort 3: Michigan/Toronto OCD Imaging Genomics Study.* Children and their parents were recruited from four academic child psychiatry sites: The Hospital for Sick Children, McMaster University, University of Michigan, and Wayne State University. Recruitment and diagnosis procedures have been described in detail elsewhere (8). All enrolled individuals had symptoms first identified before age 18. Informed consent or assent where applicable were obtained as approved by the respective institutional ethics review boards. The site clinical investigator — a child and adolescent psychiatrist — made lifetime and current axis 1 diagnoses using all sources of information according to DSM-IV criteria. OCD clinic samples (*n*=353) and controls (*n*=317) were genotyped on a variety of genotyping arrays: HumanCoreExome, PsychArray and Omni2.5. Each genotyping array was processed separately, using the same pipeline as for the Spit for Science samples. Only cases and controls were retained. Imputed data from all arrays were combined, and the association between imputed dosage and case-control status was assessed using logistic regression, using as covariates the first 3 principal components, age, sex and array identifier.

**Analyses**

**Polygenic Risk Score Prediction**

Data from IOCDF-GC/OCGAS that were used for PRS association analyses consisted of IOCDF-GC Ashkenazi Jewish (91 cases, 255 controls), IOCDF-GC European (1032 cases, 4100 controls), IOCDF-GC South African (98 cases, 157 controls) and OCGAS case/control dataset (344 cases, 1033 controls). For IOCDF-GC and OCGAS studies, the association analyses between PRS and case-control status were assessed with a logistic model, adjusted for sex and 20 ancestry dimensions from multidimensional scaling (as provided with the data); for CHOP and Michigan/Toronto, analyses were adjusted for the covariates listed above. When all studies were combined in a single PRS analysis, an additional indicator of the study was used as a covariate, and the logistic model included an interaction term between the study and the PRS to account for the between-study heterogeneity. The significance of the PRS was assessed by comparing an analysis of variance (ANOVA) of the full logistic model (with PRS and its interaction with study) to the nested model without PRS and its interaction. We used LDpred (9) to conduct the PRS calculations, which estimates a posterior mean effect size for each marker by using a Gaussian prior with point mass at zero (based on an unknown parameter ρ representing the fraction of causal markers) for the effect sizes and linkage disequilibrium (LD) information. As recommended, PRS were evaluated at the default ρ values of 1, 0.3, 0.1, 0.03, 0.01, 0.003 and 0.001, restricting to SNPs with imputation quality > 0.90, MAF>1% and present in HapMap3. We tested the association between either 1) TOCS total scores with polygenic risk for OCD case/control status at each ρ value, and 2) OCD case/control status with polygenic risk scores for TOCS scores at each ρ value. In all polygenic risk score analyses, we included genotyping array as well as PCs as covariates.

**Supplemental Results**

**OC Traits**

After sample exclusion and selection, 5018 participants were included in the GWAS analysis (out of 5645 genotyped on HumanCore and 192 genotyped on Omni. See Figure S2 for the number of samples removed during each step of QC.

The zero-inflated negative binomial distribution model was a good fit for the collapsed score (Cramer-von Mises goodness-of-fit *p=*0.83, compared to *p=*0.001 for a non-inflated negative binomial distribution, using estimated parameters).

Genetic Correlation with Other Mental Health/Medical Traits

On LD Hub, the positive correlation between TOCS total score and childhood IQ had the smallest p-value, however this correlation was not statistically significant (*r_g_*=0.64; *p*=0.19, s.e.=0.48).

**OCD Case/Control**

CHOP

After sample exclusion/selection, 406 cases and 1369 controls remained for analysis, out of a total of 421 cases and 1441 controls that were genotyped. We excluded samples because of technical quality control (*n*=11), non-European ancestry (*n*=24) and relatedness to another participant (*n*=52).

Michigan/Toronto OCD Imaging Genomics Study

A total of 690 DNA samples were genotyped on one or more arrays (HumanCoreExome *n*=45, PsychArray *n*=363, Omni2.5 *n*=282). Forty-nine samples were removed after technical exclusion; two duplicated DNAs with non-matching genome were removed; 95 samples related to or duplicates of another sample were removed; 58 samples were removed due to ancestry; and six samples were excluded due to changes in consent. After sample exclusion and sample selection, 275 cases and 205 controls remained.

**Polygenic Risk Scores**

For analyses that required individual level data from the Psychiatric Genomics Consortium (PGC), not all the data were available because of ethics approvals (IOCDFGC Dutch) and use of cases no longer included in the PGC sample (pseudo-case controls - OCGAS Trios).

**OCD Working Group of the Psychiatric Genomics Consortium Authors**

Paul D. Arnold, Kathleen D. Askland, Cristina Barlassina, O. Joseph Bienvenu, Donald Black, Michael Bloch, Helena Brentani, Beatriz Camarena, Carolina Cappi, Danielle Cath, M. Cristina Cavallini, Valentina Ciullo, David Conti, Edwin H Cook, Vladimir Coric, Bernadette A. Cullen, Danielle Cusi, Lea K. Davis, Richard Delorme, Damiaan Denys, Eske Derks, Valsamma Eapen, Christopher Edlund, Lauren Erdman, Peter Falkai, Abigail J Fyer, Daniel A. Geller, Fernando S. Goes, Hans J. Grabe, Marco A. Grados, Benjamin D. Greenberg, Edna Grünblatt, Wei Guo, Gregory L. Hanna, Ana G. Hounie, Michael Jenike, Clare L. Keenan, James L. Kennedy, Ekaterina A. Khramtsova, James A. Knowles, Janice Krasnow, Cristoph Lange, Nuria Lanzagorta, Marion Leboyer, Kung-Yee Liang, Christine Lochner, Fabio Macciardi, Brion Maher, Carol A. Mathews, Manuel Mattheisen, James T. McCracken, Nathaniel McGregor, Nicole C. McLaughlin, Euripedes c. Miguel, Benjamin Neale, Gerald Nestadt, Paul S. Nestadt, Humberto Nicolini, Erika L. Nurmi, Lisa Osiecki, John Piacentini, Danielle Posthuma, Ann e. Pulver, Steven A. Rasmussen, Scott Rauch, Margaret A. Richter, Mark A. Riddle, Stephan Ripke, Stephan Ruhrmann, Aline S. Sampaio, Jack F. Samuels, Jeremiah M. Scharf, Yin Yao Shugart, Jan Smit, Dan J. Stein, S Evelyn Stewart, Maurizio Turiel, Homero Vallada, Jeremy Veenstra-VanderWeele, Nienke Vulink, Michael Wagner, Susanne Walitza, Ying Wang, Jens Wendland, Dongmei Yu, Gwyneth Zai

**Figure Legends**

**Supplemental Figure S1: Flow Chart of Sample and SNP Exclusion for Spit for Science**

Numbers reported within square brackets are overlapping within each step. Reported non-Caucasian = not all four grandparents were parent- or self-reported to be of Caucasian European descent. HCE = HumanCoreExome array, OMNI = OMNI1 array, TOCS = Toronto Obsessive-Compulsive Scale, MAF = minor allele frequency, ASD = autism spectrum disorder, PCA = principal component analysis for population stratification, AR2 = allelic R^2^.

**Supplemental Figure S2: Principal Component Analysis Plots for Population Stratification**

Principal component (PC) analysis plots for the first three PCs showing outliers (light grey with circle with X) removed because they did not cluster with European samples (EUR, light blue). HCE = HumanCoreExome chip, omni = OMNI1 chip, AFR = African, AMR = Ad Mixed American, EAS = East Asian and SAS = South Asian.

**Supplemental Figure S3: Gene-Based Genome-Wide Analysis of OC Traits in Spit for Science**

Manhattan plot from gene-based test by MAGMA using FUMA. Four genes reached genome-wide significance (*p*=0.05/19363 protein coding genes = 2.582x10^-6^): *SH3GL2,* *PDXDC1*, *RIMBP2* and *RRN3. n*=5,018

**Supplemental Figure S4: Meta-Analysis of OCD Samples**

a) Manhattan plot and b) QQ Plot from the meta-analysis of three OCD case/control cohorts: IOCDF/OCGAS sample, CHOP and Toronto/Michigan Imaging Imaging Genomics Study (Total cases: 3,369; total controls: 8,611)

**Supplemental Figure S5: Gene-Based Genome-Wide Analysis of OCD Samples**

Manhattan plot from gene-based test by MAGMA using FUMA. No genes reached genome-wide significance (*p*=0.05/18,367 protein coding genes = 2.582x10^-6^) but *GRID2*, *DLGAP1* and *SDCBP2* approached significance (Total cases: 3,369; total controls: 8,611).
