## Supplementary figures and images for "Genome-wide Association Study of Pediatric Obsessive-Compulsive Traits: Shared Genetic Risk between Traits and Disorder"

### Suplemental Figures

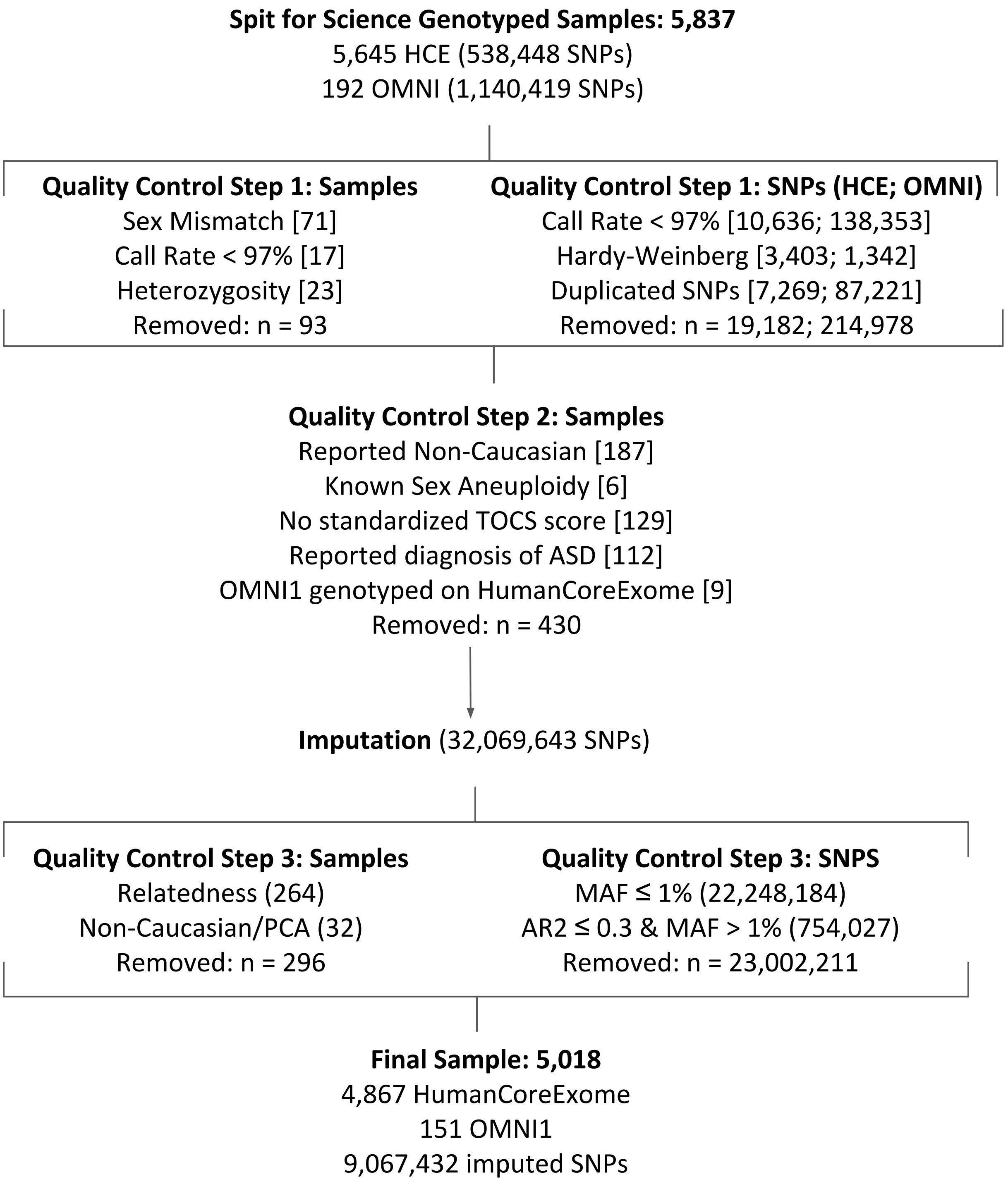

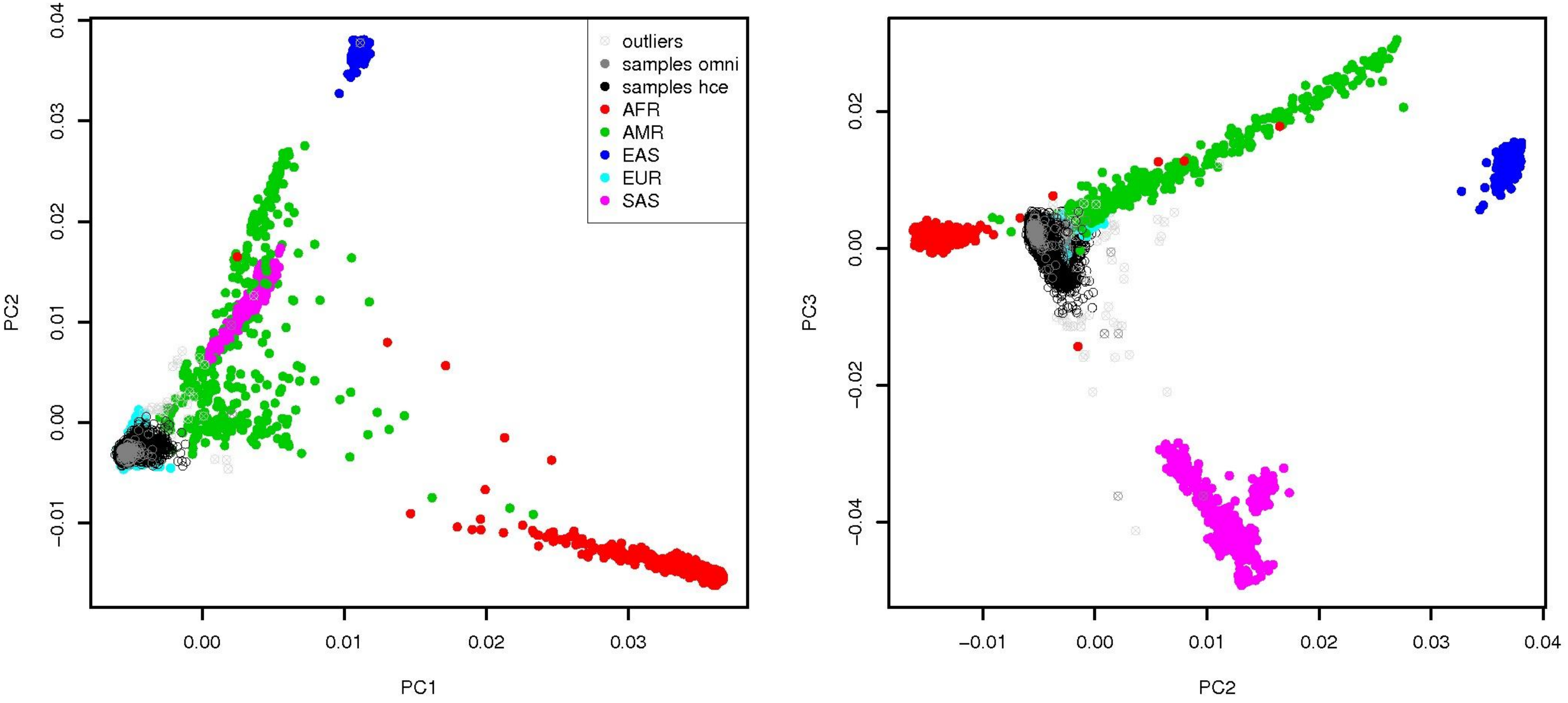

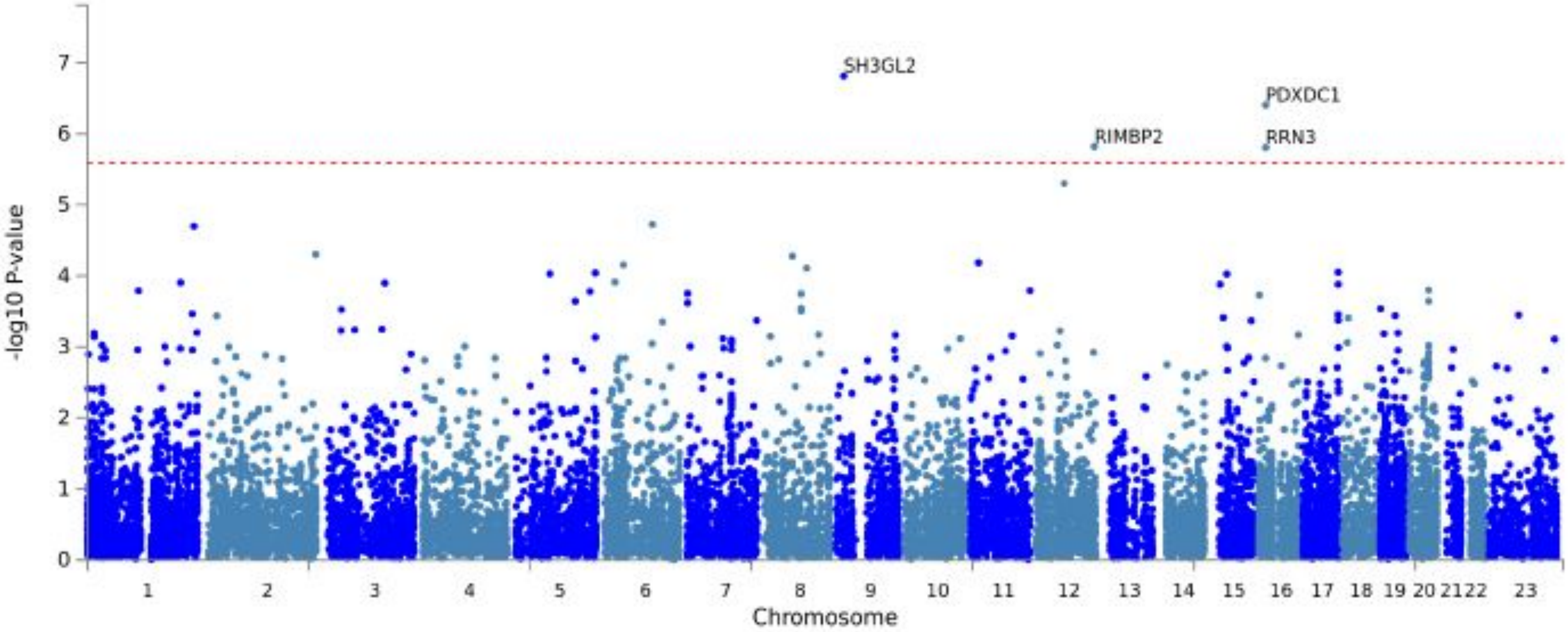

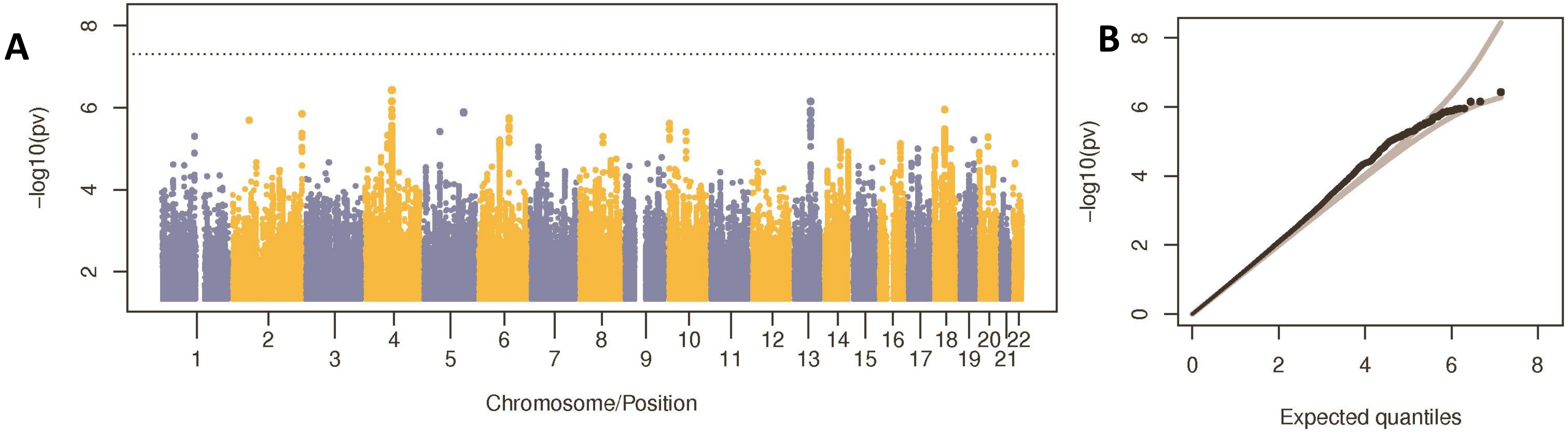

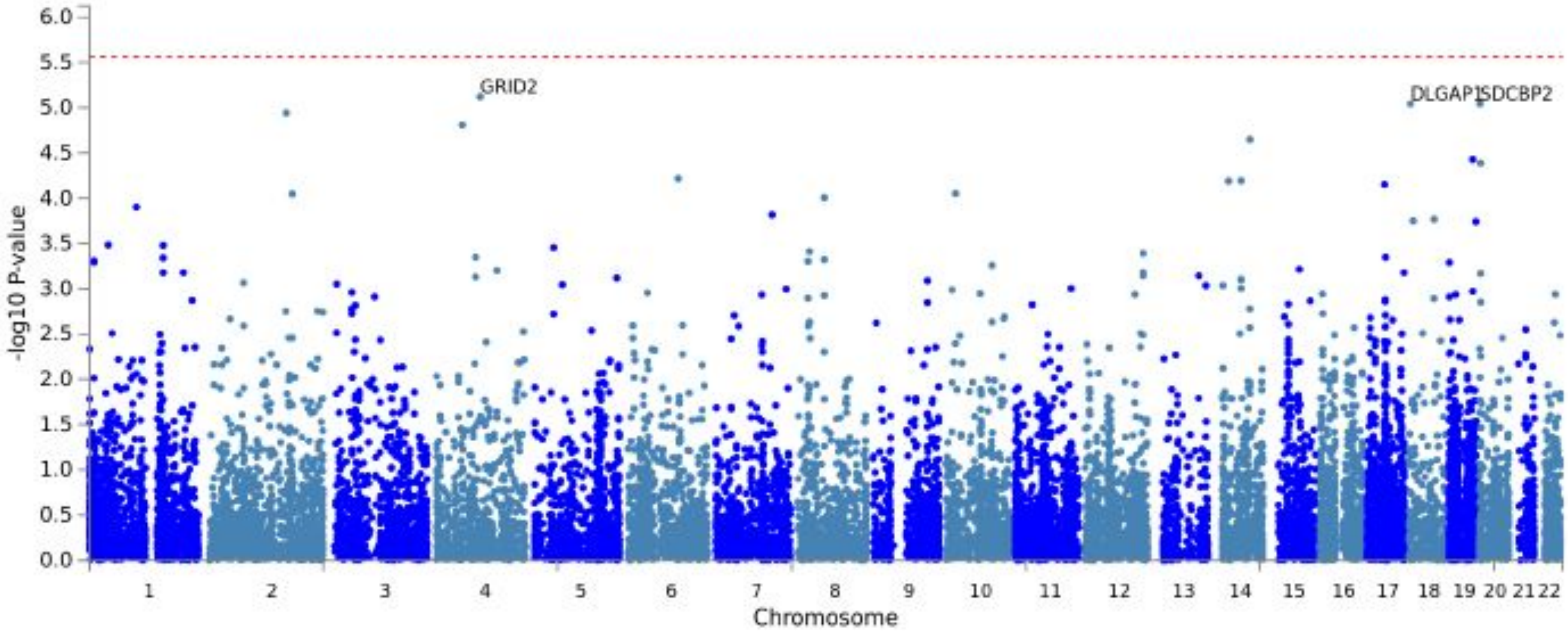
